# *Plcg2*^*M28L*^ interacts with high fat-high sugar diet to accelerate Alzheimer’s disease-relevant phenotypes in mice

**DOI:** 10.1101/2022.02.11.480032

**Authors:** Adrian L. Oblak, Kevin P. Kotredes, Ravi Pandey, Alaina M. Reagan, Cynthia Ingraham, Bridget Perkins, Chris Lloyd, Deborah Baker, Peter B. Lin, Disha M. Soni, Andy Tsai, Scott C. Persohn, Amanda A Bedwell, Kierra Eldridge, Rachael Speedy, Jill A. Meyers, Johnathon Peters, Lucas L. Figueiredo, Michael Sasner, Paul R. Territo, Stacey J. Sukoff Rizzo, Gregory W. Carter, Bruce T. Lamb, Gareth R. Howell

## Abstract

Obesity is recognized as a significant risk factor for Alzheimer’s disease (AD). Studies have supported the notion that obesity accelerates AD-related pathophysiology in mouse models of AD. The majority of studies to date have focused on the use of early-onset AD models. Here we evaluate the impact of genetic risk factors on late-onset AD (LOAD) in mice fed a high fat/high sugar diet. We focused on three mouse models created through the IU/JAX/Pitt MODEL-AD Center, LOAD1, LOAD1.*Plcg2*^*M28L*^ and LOAD1.*Mthfr*^*677C>T*^. At 2 months of age, animals were placed on a high fat/high sugar diet (HFD) that induces obesity, or a control diet (CD) that does not, until 12 months of age. Throughout the study, blood was collected to assess cholesterol and glucose. Positron emission tomography/computed tomography (PET/CT) was completed prior to sacrifice to image for glucose utilization and brain perfusion. At the completion of the study, blood and brains were collected for analysis. As expected, animals fed the HFD, regardless of genotype or sex, showed a significant increase in body weight compared to those fed the CD. Glucose and cholesterol increased as a function of HFD as well. Interestingly, LOAD1.*Plcg2*^*M28L*^ demonstrated an increase in microglia density as well as alterations in regional brain glucose and perfusion when on a HFD. These changes were not observed in LOAD1 or LOAD1.*Mthfr*^*677C>T*^ animals when fed a HFD. Furthermore, LOAD1.*Plcg2*^*M28L*^ but not LOAD1.*Mthfr*^*677C>T*^ or LOAD1 animals showed transcriptomics correlations to human AD modules. Our results show HFD affects brain health in a genotype-specific manner. Further insight into this process may have significant implications in the development of lifestyle interventions for treatment of AD.

## Introduction

Genetic and genome-wide association studies have identified variations in numerous genes that increase risk for late-onset AD (LOAD). The E4 allele of apolipoprotein E (*APOE4*) is the greatest genetic risk factor but variations in many other genes – including triggering receptor expressed on myeloid cells 2 (*TREM2*) – also contribute risk. Importantly, unlike relatively rare cases of familial AD (fAD) which are predominantly caused by mutations in *APP* or one APP processing gene, no single LOAD-associated variant is sufficient to cause AD. It is anticipated that combinations of genetic risk factors are required to develop LOAD. Studies also suggest genetic factors may only contribute between 50-70% of risk for LOAD meaning many individuals likely develop LOAD due to a combination of genetic and environmental factors[1–4]. Studies have shown that diet, obesity, cardiovascular diseases, hypertension physical activity, diabetes, educational attainment, smoking, and traumatic brain injury increase risk for LOAD and other dementias [5–8]. Some of these risk factors can be attributed to a balance between diet and exercise. For instance, a western-style diet (e.g. a diet high in fat and sugar, and low in vitamins) in combination with a sedentary lifestyle can lead to obesity, diabetes, cardiovascular disease, hypertension and increase risk for dementias[9].

The MODEL-AD (Model Organism Development and Evaluation for Late-onset AD) consortium was established to develop mouse models that more faithfully develop LOAD-relevant phenotypes compared to previous mouse models that were largely based on familial AD (fAD). Two centers, one from Indiana University (IU), The Jackson Laboratory (JAX), University of Pittsburg (Pitt) and Sage Bionetworks, and the other from University California Irvine (UCI) have focused on incorporating genetic risk factors into C57BL/6J (B6) mice and assaying human-relevant phenotypes. Initially, the IU/JAX/Pitt group created mice double homozygous for *APOE4* and *Trem2*^*R47H*^ (termed LOAD1)[10]. Mice were aged to 24 months and data showed age but not genotype was the major factor driving changes in LOAD1 mice[10]. Therefore, these mice provided a sensitized background for assessing additional genetic and environmental risk factors[10]. To further sensitize LOAD1 mice, additional LOAD genetic risk factors were added via CRISPR/CAS9. These included the M28L variant in phospholipase C Gamma 2 (*Plcg2*^*M28L*^)[11, 12] and the 677C>T variant in methylenehydrofolate reductase (*Mthfr*^*677C>T*^)[12, 13]. Transcriptomic analyses of brain tissue from LOAD1.*Plcg2*^*M28L*^ or LOAD1.*Mthfr*^*677C>T*^ mice revealed increased alignment to brain transcriptomes from human AD patients compared to B6 or LOAD1 controls [14].

Despite carrying multiple genetic risk variants for LOAD, LOAD1, LOAD1.*Plcg2*^*M28L*^ and LOAD1.*Mthfr*^*677C>T*^ did not natively recapitulate all phenotypes relevant to LOAD making these strains ideal to test the effects of environmental risk factors, such as diet. Commonly, the diet consumed by the western world is high in fat and refined sugar. This has led to a global obesity epidemic where, for instance, 42% of all Americans are considered overweight[15]. Obesity can cause a wide range of changes including increased inflammation in both the periphery and the brain [16, 17]. Inflammation is characterized in part by increased production of cytokines such as IL1β as well as activation of myeloid cells, including microglia in the brain[18]. Chronic inflammation has been shown to greatly increase risk for LOAD[19].

Multiple studies in mice have assessed the effects of a western-like diet in the context of aging and AD. For instance, one study showed diet-induced myelin breakdown in aging B6 mice that was dependent on the complement cascade[20]. Also, the addition of a HFD exacerbated AD phenotypes in mouse models relevant to early-onset AD[21–25]. In one study, Jones et al.[24] found that with HFD, male *APOE4* mice were more susceptible to metabolic disturbances, including glucose intolerance when compared to *APOE3* mice. Behavioral deficits were not observed due to the HFD, suggesting metabolic responses to HFD are dependent on both sex and APOE genotype. A second study concluded that early dysregulation of inflammation in APOE4 brains could predispose to CNS damages from various insults, including diet, and later result in the increased CNS damage normally associated with the APOE4 genotype [25]. However, the effects of a western-like diet in the context of multiple genetic risk factors for LOAD have not been studied. To address this, a commonly used HFD (high in fat and sugar, see methods) was fed to male and female LOAD1, LOAD1.*Plcg2*^*M28L*^ and LOAD1. *Mthfr*^*677C>T*^ mice from two to twelve months. Biometric measures were collected throughout the study, *in vivo* imaging to assess glucose utilization and blood perfusion in a brain region-specific manner was carried out prior to perfusion, and brain transcriptomics, neuropathological assessments and protein quantification were performed postmortem. Results showed the effects of consumption of the HFD to midlife were not uniform across all models but dependent on specific genetic risk factors. In particular, the effects of the HFD were most severe in the presence of *Plcg2*^*M28L*^.

## Results

### Body weight is increased in animals fed a high fat diet

This study consisted of three strains of animals, LOAD1, LOAD1.Mthfr^677C>T^ and LOAD1.Plcg2^M28L^ on either a normal control diet (6%) or a high fat (45%) high sugar diet. The HFD was introduced at 2 months of age in half of the animals, while the other half was maintained on the normal CD (Figure 1A). We did not find differences between males and females in either the LOAD1.Mthfr^677C>T^ or LOAD1.Plcg2^M28L^ study, therefore we collapsed male and female data together. At the beginning of the study, the average weight of the animals was 23.1 grams. Following the first month after introduction of the HFD, body weight increased significantly more in the HFD-fed mice (+8.6 ± 0.90 g) than in the CD mice (+1.3 ± 0.55 g; *P* < 0.05; Figure 1B). The weight gain continued thereafter to be progressively higher in HFD-fed mice than in mice fed CD. At the end of the study (12 mos), animals on HFD weighed on average 42.8 grams while those on the CD weighed 33.5 g (Figure 1B).

**Figure 1.**
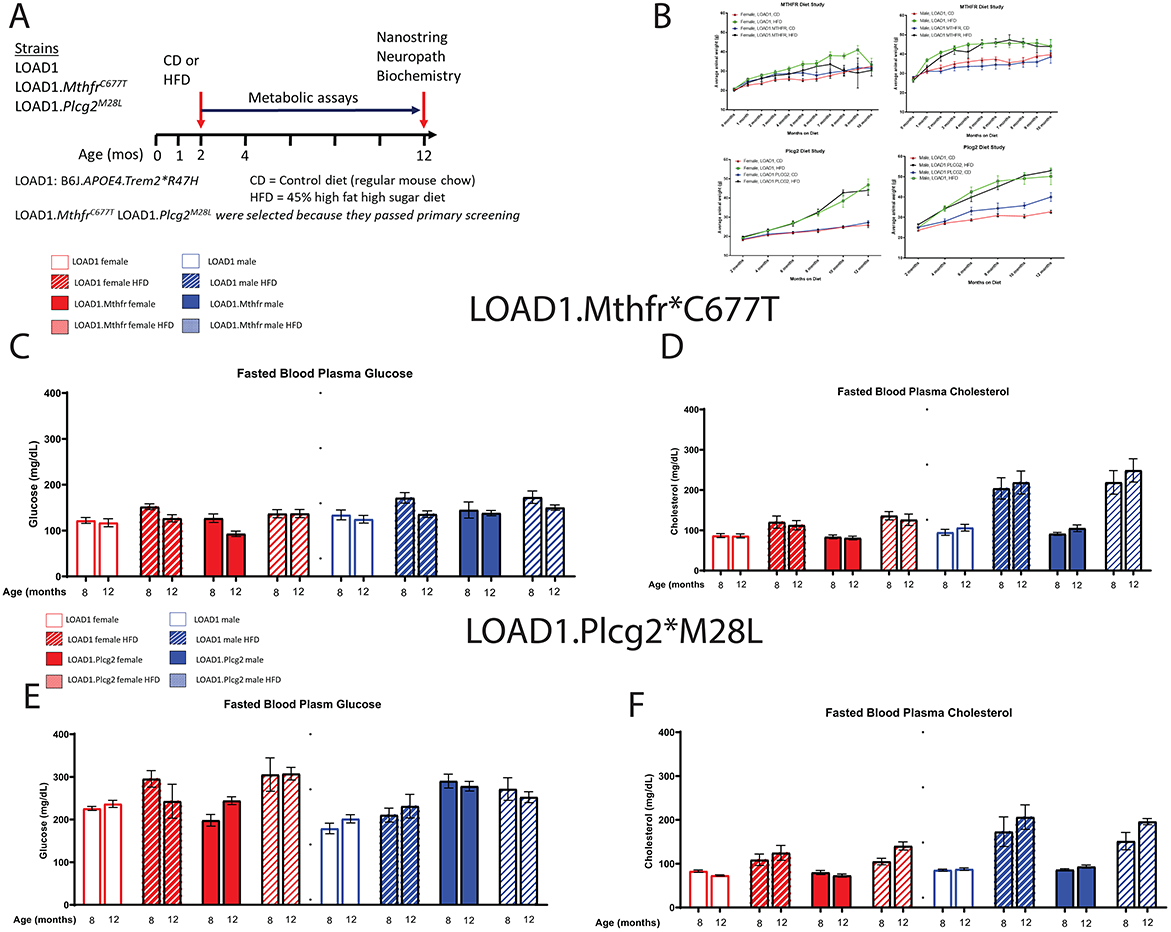
Body weight, glucose and cholesterol and increased on a high fat diet, regardless of genotype. LOAD1, LOAD1.*Mthfr*^*677C>T*^ and LOAD1.*Plcg2*^*M28L*^ mice were fed a control diet (CD) or HFD from 2 – 12 months of age (A). At multiple timepoints throughout the study, blood was collected. At the terminal time point, transcriptomics, limited neuropathology and biochemical studies were completed. Over the course of the study, regardless of genotype or sex, mice on a HFD gained more weight than animals fed a control diet (B). Glucose and cholesterol increased over time, which was related to diet, but not to genotype (C-F).

### Glucose and cholesterol are increased in animals fed a high fat diet

Bi-monthly samples were collected from fasting, non-anesthetized mice for measurements of plasma levels of glucose and cholesterol. Basal glucose was collected at 8 and 12 months in the LOAD1.*Mthfr*^*677C>T*^ study. There was no difference between the CD and HFD groups in LOAD1 and LOAD1.*Mthfr*^*677C>T*^. Similar findings in the LOAD1.Plcg2^M28L^ study were observed. At 2 mo of age, the average plasma glucose was 160.9 mg/dl and cholesterol was 81.5 mg/dl. After 10 months, the control animals in the LOAD1.*Mthfr*^*677C>T*^ study had an average 153 mg/dL glucose and 92.2 mg/dL cholesterol. The LOAD1 animals in this study had an average 162.9 mg/dL glucose and 94.9 mg/dL cholesterol.

In the LOAD1.*Plcg2*^*M28L*^ study, CD-fed animals had an average 253 mmol/l glucose (Figure 1E) and 81.2 mg/dL cholesterol (Figure 1F). The LOAD1.Plcg2^M28L^ animals fed a HFD had an average 285 mmol/l glucose and 159 mg/dL cholesterol (Figure 1E–F). In the LOAD1 control animals fed a CD fed had an average 220.7 mg/dl glucose and 81.2 mg/dL cholesterol (Figure 1E–F). The HFD-fed LOAD1 control animals had an average 271 mmol/l glucose and 179 mg/dL cholesterol (Figure 1 E–F). Data shown is comparing 8 month and 12-month-old animals (Figure 1E–F). B6 mice are considered diabetic with a fasting glucose over 240 mg/dl[26].

### Neuron number was unchanged but reactive microglia increased in LOAD1.Plcg2^M28L^ animals

A hallmark of LOAD and other dementias is the loss of neurons, particularly in the cortex and hippocampus. However, cortical and hippocampal neuronal cell loss is largely absent in most widely used mouse models of fAD and LOAD. Neuronal cell loss was also not observed in LOAD1 mice, even at 24 months of age [10]. To assess neuronal cell number in this study, we examined both the cortex and hippocampus of CD and HFD animals from both LOAD1.*Mthfr*^*677C>T*^ and LOAD1.*Plcg2*^*M28L*^ animals and controls utilizing fluorescent immunostaining (Figure 2 A–D). In both LOAD1.*Mthfr*^*C677T*^ and LOAD1.*Plcg2*^*M28L*^ animals, there was no quantitative difference in the density of neurons in either genotype when comparing the CD to the HFD (Figure 2).

**Figure 2.**
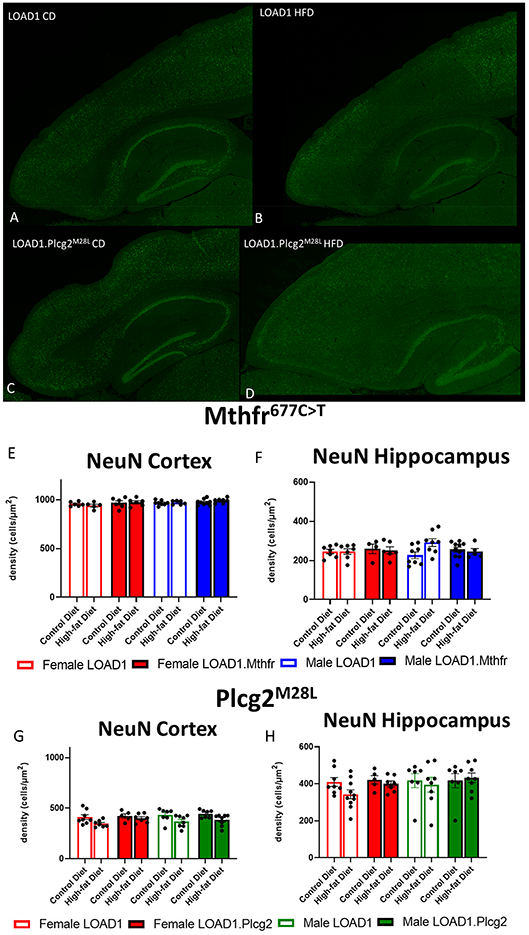
Neuron density is unaffected by a high fat diet in LOAD1.*Plcg2*^*M28L*^ and LOAD1.*Mthfr*^*677C>T*^ mice. Immunohistochemistry was completed using NeuN to visualize the density of neurons (A-D) in the LOAD1.*Plcg2*^*M28L*^ mice. HFD has no significant effect on density of neurons in either the cortex (E,G) or hippocampus (F,H) of either LOAD1.*Plcg2*^*M28L*^ or LOAD1.*Mthfr*^*677C>T*^ mice.

A second hallmark of LOAD is neuroinflammation, particularly microglia activation. In our previous study, LOAD1 mice did not show microglia activation. When analyzing the number of IBA1+ cells in the brain, compared to LOAD1 mice either on the CD or HFD we found significantly more IBA1+ cells in the cortex of female LOAD1.*Plcg2*^*M28L*^ mice fed a HFD compared to LOAD1.*Plcg2*^*M28L*^ on a CD (Figure 3). Interestingly, we did not see a difference in LOAD1.*Mthfr*^*677C>T*^ mice (Figure 3), suggesting a HFD-induced neuroinflammatory reaction that is specific to the LOAD1.*Plcg2*^*M28L*^ genotype.

**Figure 3.**
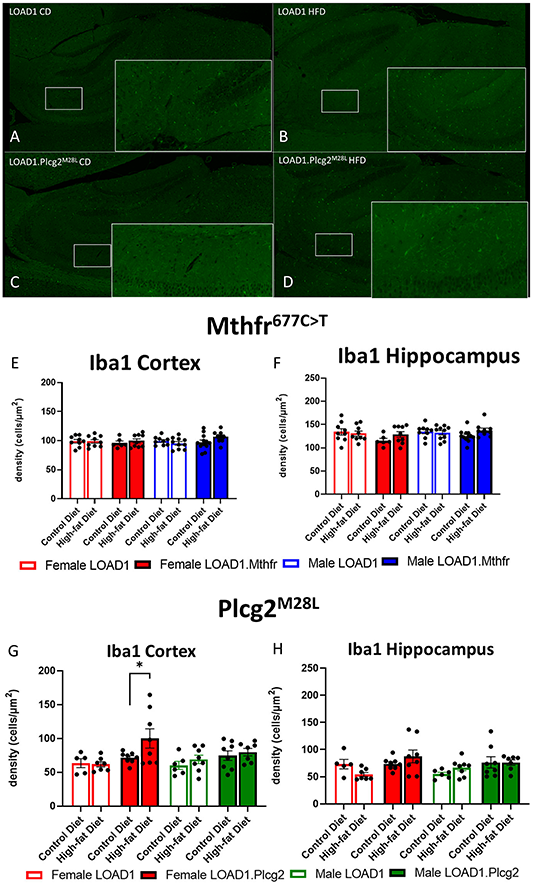
Microglial density is increased in LOAD1.*Plcg2*^*M28L*^ high fat diet mice. Immunohistochemistry was completed using Iba1 to visualize the density of microglia (A-D) in the LOAD1.*Plcg2*^*M28L*^ mice. LOAD1.*Mthfr*^*677C>T*^ mice did not show any significant changes in microglia density in the cortex (E) or hippocampus (F), regardless of diet. However, female LOAD1.*Plcg2*^*M28L*^ were found to have an increase in microglial density in the cortex of the mice fed a HFD (G). No changes were observed in LOAD1.*Plcg2*^*M28L*^ mice fed the control diet (G, H). Data suggests a gene by diet interaction that may be sex specific.

### Interaction between risk variant *Plcg2*^*M28L*^ and high fat-high sugar diet correlates with AMP-AD modules enriched for inflammatory and neuronal system associated pathways

Hallmark pathologies such as amyloid and tau accumulation and neurodegeneration have been used widely to align mouse models to human AD. However, more recently, molecular approaches have been developed by MODEL-AD[27] and others that allow more precise assessment of the relevance of mouse models to the molecular changes observed in human LOAD[28–31]. Here, we correlated the effect of each mouse perturbation (sex, HFD, genetic variants and interaction between variants and HFD) with 30 human AMP-AD co-expression modules [28]. These 30 modules, derived from different brain regions and study cohorts, were grouped into five “consensus clusters” based on similar gene content and repeated signals in the multiple regions. Both risk variants *Mthfr*^*677C>T*^ and *Plcg2*^*M28L*^ exhibited significant positive correlations (p < 0.05) with cell cycle and myelination associated modules in Consensus Cluster D and cellular stress-response associated modules in Consensus Cluster E (Figure 4A). Moreover, *Plcg2*^*M28L*^ displayed significant positive correlation (p < 0.05) with neuronal system associated modules in Consensus Cluster C (Figure 4A). Furthermore, *Plcg2*^*M28L*^ displayed significant negative correlation (p < 0.05) with immune related modules in Consensus Cluster B, while interaction between *Plcg2*^*M28L*^ and HFD displayed significant positive correlation (p < 0.05), suggesting interaction with HFD increased neuroinflammation in mice carrying the *Plcg2*^*M28L*^ risk variant (Figure 4A). In addition, interaction between *Plcg2*^*M28L*^ and HFD also exhibited significant positive correlation (p < 0.05) with extracellular matrix organization related module in Consensus Cluster A and neuronal system associated modules in Consensus Cluster C (Figure 4A). Overall, we observed AD relevant phenotypes in mice for interaction between HFD and *Plcg2*^*M28L*^ risk variant.

**Figure 4.**
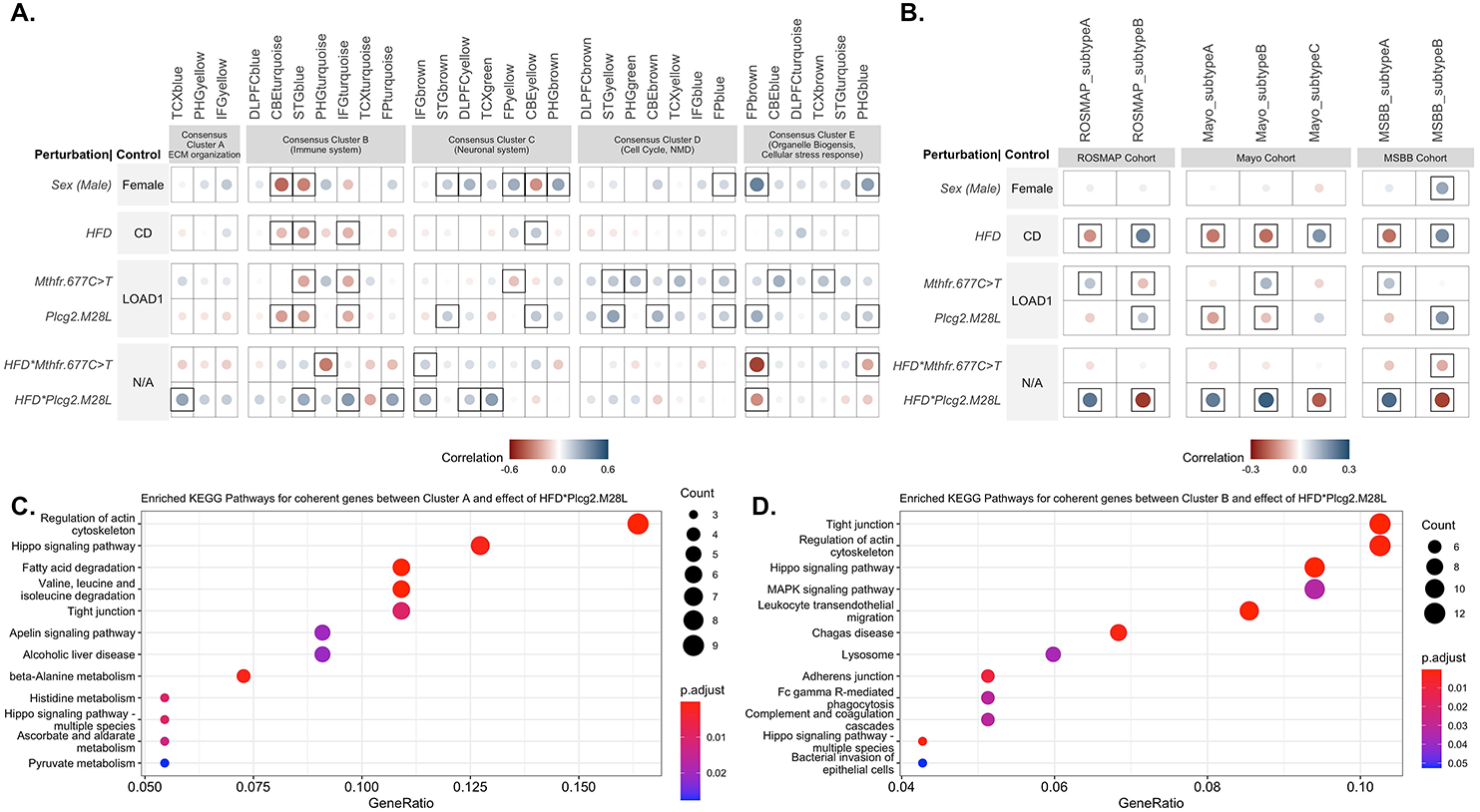
Interaction between high fat-high sugar diet and *Plcg2*^*M28L*^ in mice exhibit transcriptional changes in immune function similar to human LOAD. (A) Correlation between the effect of each mouse perturbation and 30 human co-expression modules. Note that HFD\**Plcg2.M28L* and HFD\**Mthfr.C677T* results denote interaction effects separate from the individual effects of diet and variants. Controls for corresponding rows were therefore labeled N/A as the control is not strictly defined. Circles within a square correspond to significant (*p* < 0.05) positive (blue) and negative (red) Pearson’s correlation coefficients. Color intensity and size of the circles are proportional to the Pearson’s correlation coefficient. (B) Correlation between the effect of each mouse perturbation and LOAD subtypes. The effects of interaction between HFD and *Plcg2*^*M28L*^ in mice significantly correlates with inflammatory subtype A associated with LOAD in the ROSMAP, MSBB and Mayo cohorts. Circles within a square correspond to significant (*p* < 0.05) positive (blue) and negative (red) Pearson’s correlation coefficients.(C) KEGG Pathway enrichment analysis (FDR adjusted p < 0.05) of genes exhibiting directional coherence between the effects of interaction between high fat-high sugar diet and *Plcg2*^*M28L*^ in mice and ECM organization related AMP-AD modules in Consensus Cluster A (D) KEGG Pathway enrichment analysis (FDR adjusted p < 0.05) of genes exhibiting directional coherence between the effects of interaction between HFD and *Plcg2*^*M28L*^ in mice and immune related AMP-AD modules in Consensus Cluster B.

Genes exhibiting directional coherence between the effects of interaction between HFD and *Plcg2*^*M28L*^ in mice and change in expression in AMP-AD modules in Consensus Cluster A and B, respectively were extracted. To elucidate the role of these disease-related genes, we performed KEGG pathway enrichment analysis. Genes that showed directional coherence for interaction between HFD and *Plcg2*^*M28L*^ in mice and human co-expression modules in Consensus Cluster A were enriched for “Regulation of actin cytoskeleton”, “fatty acid degradation”, and multiple “metabolism” associated pathways (Figure 4C). Genes that showed directional coherence for interaction between HFD and *Plcg2*^*M28L*^ in mice and human co-expression modules in Consensus Cluster B were enriched for multiple KEGG pathways such as “Tight junction”, “Hippo signaling pathway”, “Lysosome, and “phagocytosis” pathways (Figure 4D).

### Interaction between risk variant *Plcg2*^*M28L*^ and HFD significantly correlates with inflammatory LOAD subtypes

Next, to identify variants that resemble the inflammatory and non-inflammatory subtypes in human patients, we correlated the effect of each variant (sex, HFD, genetic variants and interaction between variants and HFD) with inflammatory and non-inflammatory subtypes associated with LOAD in the ROSMAP, MSBB and Mayo cohorts [32]. The effect of *Mthfr*^*677C>T*^ variant showed significant positive correlation (p < 0.05) with the inflammatory subtypes across all three cohorts, while effect of *Plcg2*^*M28L*^ and HFD showed significant positive correlation (p < 0.05) with the non-inflammatory subtype B in the ROSMAP and MSBB cohorts (Figure 4B). Notably, the interaction between HFD and *Plcg2*^*M28L*^ risk variant showed a strong significant positive correlation (p < 0.05) with the inflammatory subtypes across all three cohorts, while no significant correlation observed for interaction between HFD and *Mthfr*^*677C>T*^ risk variant (Figure 4B).

### Gene set enrichment analysis identified up regulation of multiple pathways of LOAD1.Plcg2^M28L^ HFD animals but not Mthfr^677C>T^

Further, Nanostring AD panel genes were ranked based on regression coefficients calculated for each factor and gene set enrichment analysis (GSEA) [33] was performed (see Methods). GSEA identified upregulation of Alzheimer’s disease, focal adhesion pathways in presence of each perturbation except *Mthfr*^*677C>T*^ (Supplementary Figure 1). Multiple immune-related pathways were upregulated and synaptic associated pathways such as GABAergic synapse, Axon guidance were downregulated in presence of HFD, and interaction between HFD and *Plcg2*^*M28L*^ (Supplementary Figure 1). Moreover, principal component analysis (PCA) using gene set enrichment score (NES) revealed HFD as a main effect in mice. We observed a clear discrimination between genetic variants with and without interaction with HFD along the first principal component (accounting for around 59% of total variation) (Supplementary Figure 3D). The second principal component account for 25% of total variance and separated *Mthfr*^*677C>T*^ and interaction between HFD and *Plcg2*^*M28L*^ factors from other factors (Supplementary Figure 1).

### Increased brain glycolysis and perfusion in *Plcg2*^*M28L*^ HFD animals

Because of the interaction between the HFD and *Plcg2*^*M28L*^ leading to a transcriptomic inflammatory signature as well as an alteration in tight junctions, it was important to determine if there were any corresponding functional deficits. To visualize alterations in regional glycolysis and perfusion, we performed *in vivo* PET/CT imaging using ^18^F-FDG (Figure 5A) and ^64^Cu-PTSM (Figure 5B), respectively. Findings were confirmed using autoradiography. Principal component analysis (PCA) comparing sex, genotype and age determined fifteen of the twenty-seven brain regions that explained 80 percent of the variance in brain glycolysis in all mice and all conditions (Figure 5 C–D). Interestingly, HFD mice show increased glucose uptake overall compared to CD mice. Utilizing PCA analysis again for blood flow analysis, thirteen of the twenty-seven brain regions explained 80 percent of the variance in blood flow in all mice and all conditions (Figure 5E–F). Both tracers were utilized in the same animals, therefore autoradiographic studies were only performed at the terminal scan (^64^Cu-PTSM; Figure 5 G–H). Interestingly, HFD mice showed increased perfusion overall compared to CD mice. Several regions that showed increased glucose metabolism and increased blood perfusion in a genotype-dependent manner are involved in memory and behavior. This data support a functional consequence of the observed interaction between HFD and *Plcg2*^*M28L*^ by IHC and transcriptomics.

**Figure 5.**
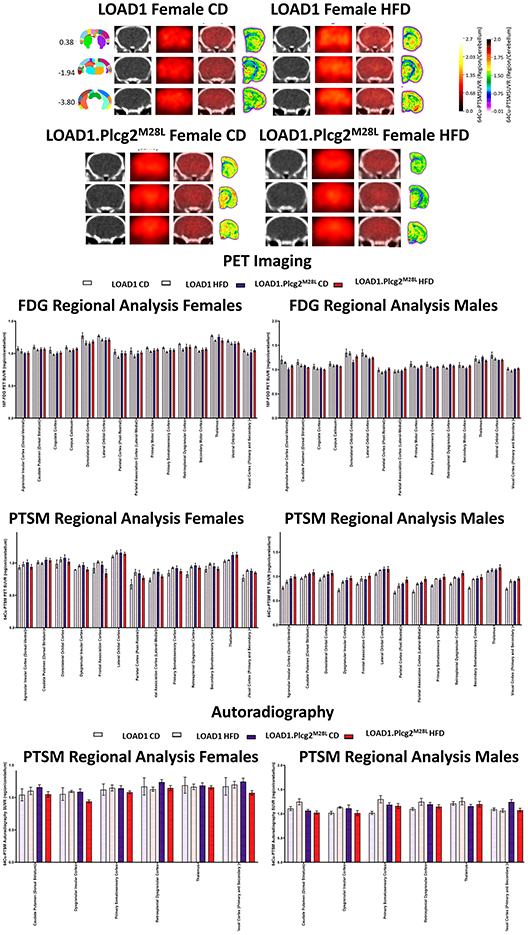
In vivo PET/CT imaging of LOAD1.Plcg2^M28L^ fed a HFD. Representative images for ^64^Cu-PTSM PET/CT and autoradiography of randomly selected 12 mo old female LOAD1.Plcg2^M28L^ mice following 10-month HFD treatment (A). In all cases, images are presented as SUVR to the cerebellum. Representative bregma image panel presented as average CT (left), PET (center-left), Fused (center-right), and Autoradiography (right). Data presented are the brain regions that explain 80 percent of the variance determined using PCA in brain glycolysis (B-C) and brain perfusion (D-E) in both females (B,D) and males (C,E). Following the terminal ^64^Cu-PTSM scans, brains were subjected to autoradiographic analysis (F-G). Data presented are the brain regions that explain 80 percent of the variance.

### Plasma cytokine and brain cytokines are altered in HFD animals

We examined the plasma and brain cytokines of mice that were fed HFD or CD (Figure 6). Consumption of a HFD leads to an increase in hormones (e.g., insulin and leptin) and cytokines in the blood that may be sensed by neurons and microglia in the brain. Based on the interaction between the LOAD1.*Plcg2*^*M28L*^ mice and the HFD, we sought to identify peripheral factors that may be driving this interaction. Analysis of cytokines in the brain revealed increases of IL-1β in HFD fed-LOAD1 animals (Figure 6A), however, the same changes were not observed in the LOAD1.*Mthfr*^*677C>T*^ animals. In addition, a significant increase in TNF-α was observed in LOAD1 males but not females (Figure 6C). No changes in IFN-γ were observed in the LOAD1. *Mthfr*^*677C>T*^ study.

**Figure 6.**
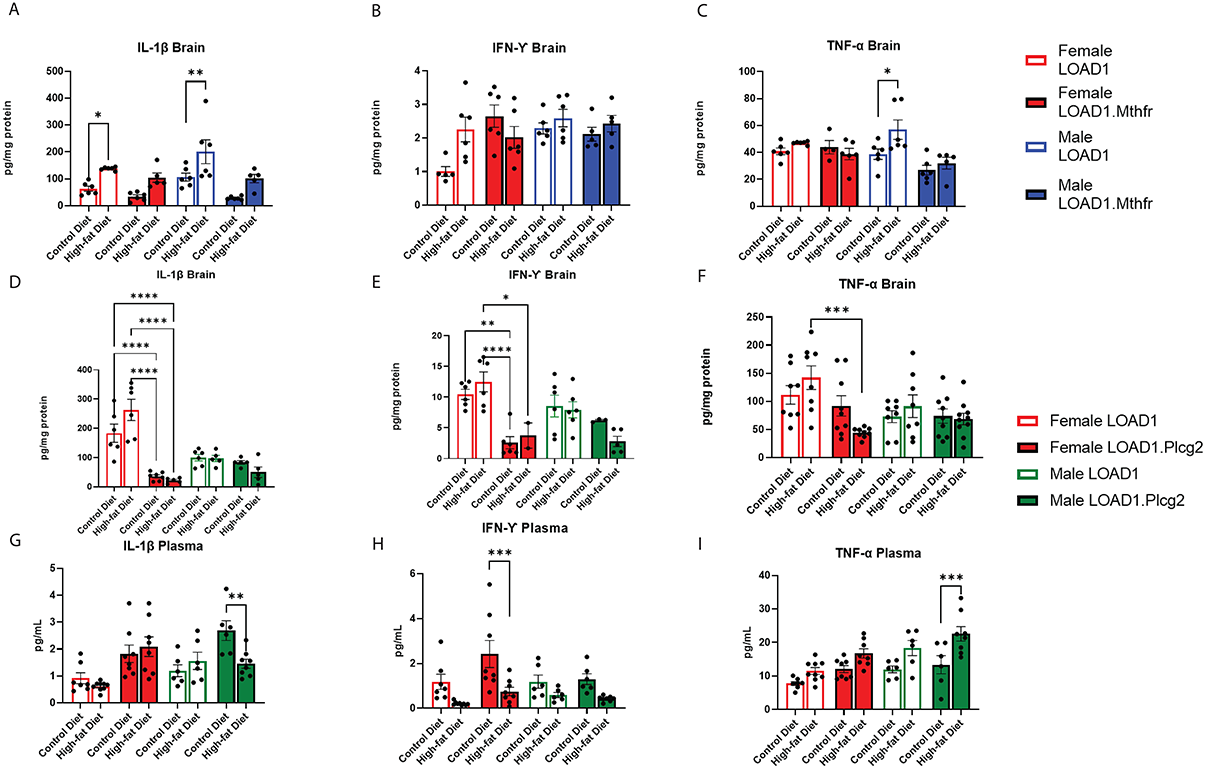
Cytokine production is altered in mice fed a HFD. In order to identify peripheral factors that may be driving the genetic by diet interaction, we examined brain (A-F) and plasma (G-I) cytokines in both LOAD1.*Plcg2*^*M28L*^ or LOAD1.*Mthfr*^*677C>T*^ mice. In LOAD1 mice, significant increases in brain IL-1β (A) and TNF-a (C) were observed in mice fed a HFD, however, LOAD1.*Mthfr*^*677C>T*^ mice did not have any significant increases. In the LOAD1.*Plcg2*^*M28L*^ mice, a significant decrease in brain IL-1β (D) and IFN-γ (E) was observed in the females, regardless of diet. The HFD did not show the same increase in brain cytokine levels as the control (LOAD1) animals, suggesting a deficiency in cytokines due to the variant. TNF-α was reduced in females as well (F), but only in the HFD group. In the plasma of LOAD1.*Plcg2*^*M28L*^ mice, we observed a reduction in IL-1β (G) and IFN-γ (H) in male and female, respectively LOAD1.*Plcg2*^*M28L*^ mice. TNF-α was elevated in the HFD LOAD1.*Plcg2*^*M28L*^ males only (I).

In LOAD1.*Plcg2*^*M28L*^, we found significant reductions in several proinflammatory cytokines (IL-1β, IFN-γ) in brain regardless of diet which were inversely correlated with plasma (Figure 6D–F). Female LOAD1 animals had an increase in plasma IL-1β in response to HFD that was not observed in males. In contrast, in the plasma of LOAD1.*Plcg2*^*M28L*^ mice, we found a significant increase in TNF-α (Figure 6I) of mice fed a HFD. However, there was a reduction in IL-1β (LOAD1.Plcg2^M28L^ males) and IFN-γ (LOAD1.Plcg2^M28L^ females), suggesting that there is a difference in peripheral and central immune responses to the HFD.

## Discussion

LOAD is a complex disorder that is caused by a combination of genetic and environmental risk factors. However, the specific combinations of genetic and environmental factors that increase risk for LOAD are not well understood. In fact, it is likely that these vary between individuals, making predictions of risk for LOAD based on their genetics and exposures imprecise. Previous studies support this, with E4FAD (APOE4+5xFAD) mice showing increased amyloid deposition, specifically more compact plaques, compared to E3FAD (APOE3+5xFAD) mice [34]. To further explore gene by diet effects, in this study, three new mouse models carrying different genetic risk factors for LOAD were exposed to a HFD for ten months (from 2-12 mos) to test the hypothesis that the reported effects of a HFD will vary depending on genetic context. The results supported our hypothesis, with mice homozygous for the *Plcg2*^*M28L*^ risk variant on a LOAD1 background showing more extreme AD phenotypes compared to either LOAD1 mice, or LOAD1 mice homozygous for *Mthfr*^*677C>T*^. To our knowledge, this is the first study to show that specific LOAD risk variants interact with a HFD to exacerbate LOAD phenotypes whereas others do not.

Diet was provided from 2 mos to 12 mos to model long-term exposure from young adult to midlife in the human population. However, our study was not designed to determine whether the changes are reversible. It is common that during an individuals’ lifetime, diet is varied, particularly as part of a weight loss program. People may go through periods of healthy eating or shorter-term extreme ‘fad’ diets such as the South Beach diet[35] or the anti-inflammatory Whole 30 diet [36]. However, although the benefits of changes in diet are observed for overt measures of health such as weight, cholesterol and blood pressure, the effects of changes in diet on the brain are not commonly tracked. Given the similarity between the effects of a HFD in humans and mice, studies where diet is varied during a lifetime, considering factors such as time on the diet and age which HFD exposure began should be performed. Here, the widely used 45% HFD (high fat, high sugar diet, ResearchDiet® feed D12451i) was used to mimic diet-induced obesity (DIO). This allowed the effects of DIO to be determined within the 12 months of this study and further studies would be needed to assess longer term effects to old age. Also, the HFD used here is somewhat extreme – not many individuals consume 45% fat daily. Therefore, to further align mouse studies to the human situation, alternative diets should be tested such as a recently developed western diet that incorporates 16% fat that was designed to model the more common diet consumed in the western world [37, 38].

The interaction between *Plcg2*^*M28L*^ and a HFD was determined using the Nanostring Mouse AD panel. These analyses indicated Plcg2^M28L^:HFD (interaction) resulted in changes in the expression of genes enriched for inflammatory pathways. The direction of the changes correlated with gene expression changes observed in human AD based on the AMP-AD modules [14].

Many of these ‘inflammation’ genes are expressed by microglia including *Csf1*, *Tgfbr2*, *Fcer1g*, *Trem2*, and *Tyrobp*. PLCG2 is a critical signaling element for various immune receptors and is a key regulatory hub gene for immune signaling. PLCG2 is expected to be important in AD due to the previous findings that a hypermorphic variant in PLCG2, rs72824905, is protective against AD risk. However, the role of PLCG2 has not yet been comprehensively explored [11, 39]. A previous study has reported that reduced PLCG2 gene expression alter microglial phenotypes in 5XFAD mice, affect plaque pathology, and drive distinct transcriptional phenotypes of microglia in the presence of amyloid pathology[11]. The HFD used here is reported to create a more inflammatory environment in the brain, causing microglia activation. Previous HFD studies have been associated with increased microglial activity in wild-type mice [40, 41]. In the present study, transcriptomic changes were observed in the neuroinflammation module, indicative of alterations in microglial function. Consistent with the transcriptomic changes, this study revealed an increase in microgliosis in the hippocampus of LOAD1.Plcg2^M28L^ mice, but not in LOAD1.*Mthfr*^*677C>T*^.

Recent *in vivo* imaging studies have suggested that microglia displayed higher glucose uptake than neurons and astrocytes [42]. In Alzheimer’s Disease patients, glucose uptake was measured using FDG-PET and increases in glucose uptake were observed with an increase in microglial activity (utilizing TSPO-PET)[42]. In the present study we observed alterations in FDG activity in multiple brain areas in LOAD1 mice carrying the *Plcg2*^*M28L*^ variant. It is possible that the predicted loss of function of this variant alters the microglial state, leading to a dampened FDG signal and reduced function, which would agree with the reduced cytokine release that was observed in the brain tissue. Although we observed increases in microglial number in female LOAD1.*Plcg2*^*M28L*^ mice, this was only in the hippocampus of HFD animals, suggesting a potential compensatory effect. The present study suggests that FDG-PET may provide evidence of microglial state *in vivo*.

Interestingly, the interaction between *Mthfr*^*677C>T*^ and HFD did not result in gene expression changes relating to inflammation and therefore PET/CT was not performed in LOAD1.*Mthfr*^*677C>T*^ mice. MTHFR is a key enzyme in the folate/methionine/homocysteine pathway. Variations in MTHFR, particularly the *Mthfr*^*677C>T*^ variant, are commonly associated with cardiovascular diseases, Alzheimer’s disease, and vascular dementia. In humans, *Mthfr*^*677C>T*^ reduces liver enzyme activity resulting in a decrease in enzyme function. Common effects of this include increases in homocysteine which is reported to increase risk for vascular inflammation and dysregulation. MTHFR is also expressed in multiple cells in the brain particularly in vascular-related cells such as endothelial and vascular smooth muscle cells. Supporting a role for MTHFR in cerebrovascular function, B6 mice homozygous or heterozygous for *Mthfr*^*677C>T*^ show reduced enzyme activity in both the liver and brain, elevated levels of homocysteine, cerebral blood flow deficits, reduced collagen 4 in the brain and neurovascular damage by electron microscopy[43]. However, unlike *Plcg2*^*M28L*^, a function for MTHFR in microglia has not been reported, further suggesting the interaction between *Plcg2*^*M28L*^ and HFD is driven by changes in microglial function.

Despite the increased alignment at the gene expression of LOAD1.*Plcg2*^*M28L*^ on a HFD to human LOAD compared to strain-matched mice fed a CD, LOAD1.*Plcg2*^*M28L*^ mice fed a HFD still lack amyloid-beta plaques. This is not surprising as mice require mutations or engineering of the human sequence to drive plaque deposition neither of which is present in these mice. Notably this allows us to study the effect of LOAD genetic risk x aging in the absence of amyloid pathology. Also, neuronal loss was not detected in LOAD1.*Plcg2*^*M28L*^ mice on a HFD – at least in the brain regions assessed up to 12 months of age – suggesting additional pathways/processes as well as increased aging need to be perturbed to stress neurons to the point of death. The presence of age-dependent amyloid accumulation would be expected to further modify LOAD phenotypes and possibly result in neuronal cell loss. Therefore, a humanized amyloid-beta (hAβ) allele has been created by MODEL-AD and added to LOAD1 (to create LOAD2; B6.*APOE4*.*Trem2^R47H^*.hAβ) as well as to LOAD1.*Plcg2*^*M28L*^ and LOAD1.*Mthfr*^*677C>T*^ to create LOAD2.*Plcg2*^*M28L*^ and LOAD2.*Mthfr*^*C677T*^ respectively. Male and female test and control mice are being provided a HFD and being aged to 18-24 months to determine the long-term effects of combining *APOE4*, *Trem2^R47H^*, *Plcg2*^*M28L*^ or *Mthfr*^*677C>T*^ risk variants in combination with aging and the more amyloidogenic humanized hAβ.

Hippocampal injury is found in HFD-fed animals in response to increased blood-brain barrier permeability [44] due to circulating proinflammatory adipokines and reactive glial cytokine production [45]. Increased blood-brain barrier permeability that allows proinflammatory proteins into the brain can initiate neuroinflammation and stimulate neurodegeneration [46]. Peripheral cytokines can act on the brain to trigger cytokine production [47]. In the present study, we also observed increases in tissue perfusion utilizing ^64^Cu-PTSM that could be related to an increase in permeability of the blood brain barrier, which would provide a means for peripheral cytokines to more easily enter the central nervous system. Additionally, data in animals revealed that insensitivity of the leptin receptor [48] upregulated the transcriptional activity of interleukin-6 (IL-6), TNF-α, and IL-1β in the hippocampus compared to control [49]. In the present study, we found alterations in several peripheral cytokines including IL-1β, IFN-γ and TNF-α in only the LOAD1.*Plcg2*^*M28L*^ animals. In the LOAD1 and LOAD1. *Mthfr*^*677C>T*^ animals, TNF-α and IL-1β were upregulated in LOAD1 animals fed a high fat diet, but not the LOAD1.*Plcg2*^*M28L*^ animals. However, TNF-α, IL-1β, and IFN-γ, were downregulated in LOAD1.*Plcg2*^*M28L*^ female brains, regardless of diet. Taken together, these data suggest that consumption of HFD produces more proinflammatory cytokines to enter the brain and initiate an inflammatory cascade, which leads to transcriptional changes that correlate with human LOAD.

Despite the fact that all mice carried the *APOE4* and *Trem2*^*R47H*^ variants, linear modeling identified the specific effects of the *Plcg2*^*M28L*^ and *Mthfr*^*677C>T*^ variants as well as the *Plcg2*^*M28L*^:HFD and *Mthfr*^*677C>T*^:HFD interactions. However, data now suggest that although APOE4 and *TREM2*^*R47H*^ are strong genetic risk factors for LOAD in humans, *TREM2*^*R47H*^ may be reducing the effect of the APOE4 variant when present together[10]. This may be due in part to the reduced expression levels of *Trem2* in *Trem2*^*R47H*^ mice caused by a cryptic splice site that results in an aberrant splice form (JAX#27918). MODEL-AD has created a new *Trem2^R47H^* allele (*Trem2*R47H ^HSS^*) that incorporates a human splice site (HSS) and restores *Trem2* expression to normal levels – at least in young wild type B6 mice (JAX#33781). Future studies will incorporate APOE4 or *Trem2*R47H*^*HSS*^ in combination with the hAβ allele and a recently created humanized MAPT-GR (JAX # 35398 and 33668) to study the interaction between HFD and Tau pathology in the context of LOAD. These mice express the human MAPT (H1 or H2 haplotype, respectively) and MAPT-AS1 transcripts and the typical MAPT protein isoforms.

In summary, combining genotype and environmental factors (e.g. HFD), may lead to better translational and preclinical models of LOAD. The interactions we have observed here that correlate with LOAD in the human population suggest that the effects of a HFD are genotype-specific and further investigation is needed to resolve the mechanistic interactions between genetics and diet.

### Data Availability Statement

The original contributions presented in the study are publicly available. The molecular, bioinformatics and imaging data are available via the AD Knowledge Portal: https://adknowledgeportal.org (permission was obtained for this material through a Creative Commons CC-BY license). The data, analyses, and tools are shared early in the research cycle without a publication embargo on secondary use. Data are available for general research use according to the following requirements for data access and data attribution (https://adknowledgeportal.org/DataAccess/Instructions).

## Methods

### Animal Housing Conditions at Indiana University and The Jackson Laboratory

All animals were created and obtained from The Jackson Laboratory (JAX). Models used here are congenic to the C57BL/6J (JAX# 000664) (B6) strain. LOAD1 is homozygous for both *APOE4 and Trem2*^*R47H*^ (JAX ID:28709). B6.*Plcg2*^*M28L*^/*APOE4*/*Trem2*^*R47H*^ (triple homozygous, LOAD1.*Plcg2^M28L^,* JAX ID:30674) was created using CRISPR/Cas9 to introduce the M28L LOAD risk variant into LOAD1 mice. B6.*Mthfr*^*677C>T*^/*APOE4*/*Trem2*^*R47H*^ (triple homozygous, LOAD1.*Mthfr*^*677C>T*^, JAX ID:30922), was also created using CRISPR/Cas9 to introduce the 677TC>T variant in to LOAD1 mice[43]. More details on strain creation are provided on the AD Knowledge Portal.

The effects of HFD on LOAD1.*Plcg2*^*M28L*^ mice were assessed at Indiana University (IU), with LOAD1.*Mthfr*^*677C>T*^ mice assessed at JAX. LOAD1 mice acted as site-matched controls. An additional cohort of B6 mice were assessed at IU as a strain control. At IU, for experimental cohorts, LOAD1.*Plcg2*^*M28L/+*^ mice were intercrossed to create LOAD1.*Plcg2*^*M28L*^ triple homozygous and LOAD1 litter-matched control mice. At JAX, for experimental cohorts, LOAD1.*Mthfr*^*677TC>T/+*^ mice were intercrossed to create LOAD1.*Mthfr*^*677TC>T*^ triple homozygous and LOAD1 litter-matched control mice. Up to five mice were housed per cage with SaniChip bedding and provided LabDiet® 5K52/5K67 (6% fat, control diet, CD). Mouse rooms were kept on a 12:12 light:dark schedule with the lights on from 7:00 am (6:00 am at JAX) to 7:00 pm daily (6:00 pm at JAX). Mice were initially ear-punched for identification and then following genotype confirmations microchipped using p-chip system (PharmaSeq) microchips placed at the base of the tail. At 2 months of age (mos), experimental cohorts were randomized into two groups: Group 1 continued on CD ad libitum and group 2 was provided ad libitum ResearchDiet® feed D12451i (45% high fat; 35% carbohydrates, HFD). All procedures were approved by the Institutional Animal Care and Use Committees (IACUC) at IU and JAX. Where possible, all housing and procedures were standardized and aligned across sites. Unless specified, n=10 for each sex, genotype and diet were used.

### Perfusion and Preparation of Blood and Tissue Samples

Mice were anesthetized to the surgical plane of anesthesia with tribromoethanol at 12 months of age. Under complete anesthesia, animals were euthanized by decapitation and perfused through the heart with ice-cold PBS. Blood and brain tissue were collected immediately after euthanasia. Trunk blood was centrifuged for 15-20 min at 4°C at 14,500 RPM, and the plasma was stored at −80°C.

### Blood Plasma Analysis

Blood was collected longitudinally from non-fasted mice aged 2, 4, 6, 8, 10 or 12 months via a cheek puncture. For terminal timepoint collections, mice were anesthetized, and blood was extracted by left ventricle cardiac puncture with a 25 g EDTA-coated needle before PBS perfusion. Approximately 500 μL of whole blood was transferred to a MAP-K2 EDTA Microcontainer (BD, Franklin Lakes, NJ) on ice and centrifuged at 4°C at 4388 x g in a pre-chilled ultracentrifuge for 15 min. Without disturbing the red blood cell fraction, serum supernatant was pipetted into a chilled cryovial with a p200 tip and immediately snap-frozen on dry ice for 10 min. Samples were stored at −80°C and later thawed for analysis of glucose, total cholesterol, low-density lipoprotein (LDL), high-density lipoprotein (HDL), triglyceride, and non-essential fatty acid (NEFA) levels with the Siemens Advia 120 (Germany).

### Histology and Fluorescent Immunostaining

After perfusion and dissection, the left side of each mouse brain was fixed in 4% paraformaldehyde. After a transfer to 10% sucrose the following day, the brains were transferred to 30% sucrose for storage. Brains were sectioned, coronally and sagittally, at 10-20 µm on a freezing microtome.

#### NeuN Staining

Sections were washed and blocked for one hour in 10% host goat serum. Sections were incubated overnight at 4°C in a solution containing the antibodies NeuN rabbit (ab104225, 1:1000, Abcam). After additional washes, sections were incubated for an hour at room temperature in secondary solution containing the fluorescent markers goat anti-rabbit 488 (A11034, 1:1000, Invitrogen). After one additional wash, sections were mounted on charged slides, counterstained, and coverslipped with Prolong Gold Antifade Mountant with DAPI.

#### Iba1 Staining

Sections were washed and then blocked with 10% host goat serum for 2 hours. Sections were incubated overnight at 4°C in a solution containing the Recombinant Anti-Iba1 antibody rabbit (ab178847, 1:500, abcam). After additional washes, sections were incubated for an hour at room temperature in secondary solution containing the fluorescent markers goat anti-rabbit 488 (A11034, 1:1000, Invitrogen). After one additional wash, sections were mounted on charged slides, counterstained, and coverslipped with Prolong Gold Antifade Mountant with DAPI.

Microscopy was used to view and capture images of the immunofluorescent stains with a Leica DM6 B and DFC7000 GT camera using Leica Microsystems’ LAS X software. Further image capture and image scanning was performed on an Andor Zyla 5.5 sCMOS camera with an Aperio Versa scanner and Versa software.

### RNA Preparation and Correlation Analysis

Similar to the methods previously described[27, 50], total RNA was extracted from frozen right brain hemispheres using the MagMAX mirVana Total RNA Isolation Kit (ThermoFisher) and the KingFisher Flex purification system (ThermoFisher, Waltham, MA; n=6 per sex/genotype/diet). RNA concentration and quality were assessed using the Nanodrop 2000 spectrophotometer (Thermo Scientific) and the RNA Total RNA Nano assay (Agilent Technologies, Santa Clara, CA). The NanoString Mouse AD gene expression panel was used for gene expression profiling on the nCounter platform (NanoString, Seattle, WA) as described by the manufacturer. nSolver software was used for generating raw NanoString gene expression values. NanoString data was normalized by dividing raw counts within a lane by geometric mean of the housekeeping genes from the same lane[14]. Next, normalized count values were log-transformed for downstream analysis.

To determine the effects of each variant, we fit a multiple regression model as [51]:

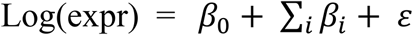

Where i denotes the sex (male), high fat diet, genetic variants (*Mthfr*^*677C>T*^, *Plcg2*^*M28L*^), and interaction terms between high fat diet and each variant (HFD* *Mthfr*^*677C>T*^, HFD* *Plcg2*^*M28L*^), and log(expr) represents log-transformed normalized count of Nanostring gene expression panel [27]. In this formulation, the standard chow diet and LOAD1 genetic background serve as controls.

Next, we have computed Pearson correlation between gene expression changes (log fold change) in human AD cases versus controls and the effect of each mouse perturbation for each gene in Nanostring panel [27, 51] using cor.test function built in R as:

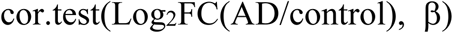

from which we obtained both the correlation coefficient and the significance level (p-value) of the correlation. Log2FC values for human transcripts were obtained through the AD Knowledge Portal [52](https://www.synapse.org/#!Synapse:syn14237651).

### KEGG Pathway enrichment analysis

KEGG pathway enrichment analysis were performed using clusterprofiler package [53] within the R software environment. Pathways were determined to be significant after multiple testing correction (FDR adjusted p < 0.05).

### In vivo PET/CT Imaging

To assess regional glycolysis and tissue perfusion, mice were non-invasively imaged via PET/CT (n=10 mice/sex/genotype/age). To measure regional blood flow, copper-pyruvaldehyde-bis(N4-methylthiosemicarbazone) (^64^Cu-PTSM)[54], which has a very high first pass (>75%) extraction[55], and glutathione reductase redox trapping of copper[55], was administered via tail vein in awake subjects and was given a 2 min uptake period prior to imaging. To measure regional glycolytic metabolism, 2-fluoro-2-deoxyglucose (^18^F-FDG) was administered via tail vein in awake subjects and given a 30 min uptake period prior to imaging. Post uptake, mice were induced with 5% isoflurane (95% medical oxygen) and maintained during acquisition with 1-2% isoflurane at 37°C. To provide both anatomical structure and function, PET/CT imaging was performed with a Molecubes β-X-CUBE system (Molecubes NV, Gent Belgium). For PET determination of blood flow and metabolism, calibrated listmode PET images were acquired on the β-CUBE and reconstructed into a single-static image using ordered subset expectation maximization (OSEM) with 30 iterations and 3 subsets[56]. To provide anatomical reference, and attenuation maps necessary to obtain fully corrected quantitative PET images, helical CT images were acquired with tube voltage of 50 kV, 100 mA, 100 μm slice thickness, 75 ms exposure, and 100 μm resolution. In all cases, images were corrected for radionuclide decay, tissue attenuation, detector dead-time loss, and photon scatter according to the manufacturer’s methods[56]. Post-acquisition, all PET and CT images were co-registered using a mutual information-based normalized entropy algorithm[57] with 12 degrees of freedom and mapped to stereotactic mouse brain coordinates[58]. Finally, to quantify regional changes, voxels of interest (VOI) for 27 brain (54 bilateral) regions were extracted and analyzed for SUVR according to published methods[59].

#### Autoradiography

To provide secondary confirmation of the *in vivo* PET images, and to quantify tracer uptake regionally, brains were extracted post rapid decapitation, gross sectioned along the midline, slowly frozen on dry ice, then embedded in cryomolds with Optimal Cutting Temperature (OCT) compound (Tissue-Tek). Thin frozen sections (20 um) were obtained via cryotomy at prescribed bregma targets (n=6 bregma/mouse, 6 replicates/bregma) according to stereotactic mouse brain coordinates [60]. Sections were mounted on glass slides, air dried, and exposed on BAS Storage Phosphor Screens (SR 2040 E, Cytiva Inc.) for up to 12 hrs. Post-exposure, screen were imaged via Typhoon FL 7000IP (GE Medical Systems) phosphor-imager at 25 μm resolution along with custom ^18^F or ^64^Cu standards described previously [61].

#### Image Analysis

All PET and MRI images were co-registered using a ridged-body mutual information-based normalized entropy algorithm [62] with 12 degrees of freedom, and mapped to stereotactic mouse brain coordinates [60] using MIM 7.0.5 (MIM Software Inc., Beachwood, Ohio). Post-registration, 56 regions bilateral regions were extracted via brain atlas, and averaged to yield 27 unique volumes of interest that map to key cognitive and motor centers that includes: Agranular Insular Cortex; Auditory Cortex; Caudate Putamen, Cerebellum; Cingulate Cortex; Corpus Callosum; Dorsolateral Orbital Cortex; Dorsintermed Entorhinal Cortex; Dysgranular Insular Cortex; Ectorhinal Cortex; Fornix; Frontal Association Cortex; Hippocampus; Lateral Orbital Cortex; Medial Orbital Cortex; Parietal Cortex; Parietal Association Cortex; Perirhinal Cortex; Prelimbic Cortex; Primary Motor Cortex; Primary Somatosensory Cortex; Retrosplenial Dysgranular Cortex; Secondary Motor Cortex; Secondary Somatosensory Cortex; Temporal Association Cortex, Thalamus; Ventral Orbital Cortex; Visual Cortex. For autoradiographic analysis, tracer uptake was quantified on hemi-coronal sections by manually drawing regions of interest for 17 regions of interest (i.e. Auditory Cortex, Caudate Putamen, Cerebellum, Cingulate Cortex, Corpus Callosum, Dorso-intermed Entorhinal Cortex, Dysgranular Insular Cortex, Ectorhinal Cortex, Hippocampus, Hypothalamus, Medial Septum, Primary Motor Cortex, Primary Somatosensory Cortex, Retrosplenial Dysgranular Cortex, Temporal Association Cortex, Thalamus, Visual Cortex) on calibrated phosphor screen at bregma 0.38, −1.94, and −3.8 mm using MCID (InterFocus Ltd). To permit dose and brain uptake normalization, Standardized Uptake Value Ratios (SUVR) relative to the cerebellum were computed for PET and autoradiograms for each subject, genotype, and age as follows:

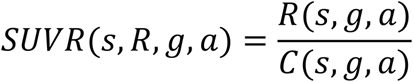

where, *s*, *g*, *a*, *R*, and *C* are the subject, genotype, age, region/volume of interest, cerebellum region/volume of interest. In all cases, region/volumes of interest were subjected to PCA analysis, and regions which explain the top 80% of the variance were analyzed for differences with time and genotype using a Two-Way ANOVA (Prism, GraphPad Inc.), where significance was taken at p<0.05.

### Homogenization and Protein Extraction

Each hemibrain was weighed prior to homogenizing in tissue protein extraction reagent (T-PER ThermoScientific; 1mL per 100mg of tissue weight) supplemented with protease and phosphatase inhibitors cocktail (Sigma-Aldrich). Total protein concentration was measured using bicinchoninic acid (BCA; Pierce). Hemibrain lysates were then aliquoted and kept in the −80°C freezer for long-term storage. The primary supernatant was utilized to analyze the content of proinflammatory cytokines.

### Cytokine Panel Assay

Mouse hemibrain samples were assayed in duplicate using the MSD Proinflammatory Panel I (K15048D; MesoScale Discovery, Gaithersburg, MD, USA), a highly sensitive multiplex enzyme-linked immunosorbent assay (ELISA). This panel quantifies the following 10 proinflammatory cytokines in a single small sample volume (25 μL) of supernatant using an electrochemiluminescent detection method (MSD): interferon γ (IFN-γ), interleukin (IL)-1β, IL-2, IL-4, IL-6, IL-8, IL-10, IL-12p70, IL-13, and tumor necrosis factor α (TNFα). The mean intra-assay coefficient for each cytokine was <8.5%, based on the cytokine standards. Any value below the lower limit of detection (LLOD) for the cytokine assay was replaced with ½ LLOD of the assay for statistical analysis.

## Acknowledgements

The MODEL-AD Center is supported through funding by NIA grant U54AG054345. The results published here are in whole or in part based on data obtained from the AD Knowledge Portal (https://adknowledgeportal.org). Study data were provided by the Rush Alzheimer’s Disease Center, Rush University Medical Center, Chicago. Data collection was supported through funding by NIA grants P30AG10161 (ROS), R01AG15819 (ROSMAP; genomics and RNAseq), R01AG17917 (MAP), R01AG36836 (RNAseq), the Illinois Department of Public Health (ROSMAP), and the Translational Genomics Research Institute (genomic). Additional phenotypic data can be requested at www.radc.rush.edu. Mount Sinai Brain Bank data were generated from postmortem brain tissue collected through the Mount Sinai VA Medical Center Brain Bank and were provided by Dr. Eric Schadt from Mount Sinai School of Medicine. The Mayo RNAseq study data was led by Dr. Nilüfer Ertekin-Taner, Mayo Clinic, Jacksonville, FL as part of the multi-PI U01 AG046139 (MPIs Golde, Ertekin-Taner, Younkin, Price). Samples were provided from the following sources: The Mayo Clinic Brain Bank. Data collection was supported through funding by NIA grants P50 AG016574, R01 AG032990, U01 AG046139, R01 AG018023, U01 AG006576, U01 AG006786, R01 AG025711, R01 AG017216, R01 AG003949, NINDS grant R01 NS080820, CurePSP Foundation, and support from Mayo Foundation. Study data includes samples collected through the Sun Health Research Institute Brain and Body Donation Program of Sun City, Arizona. The Brain and Body Donation Program is supported by the National Institute of Neurological Disorders and Stroke (U24 NS072026 National Brain and Tissue Resource for Parkinsons Disease and Related Disorders), the National Institute on Aging (P30 AG19610 Arizona Alzheimers Disease Core Center), the Arizona Department of Health Services (contract 211002, Arizona Alzheimers Research Center), the Arizona Biomedical Research Commission (contracts 4001, 0011, 05-901 and 1001 to the Arizona Parkinson’s Disease Consortium) and the Michael J. Fox Foundation for Parkinsons Research.

---

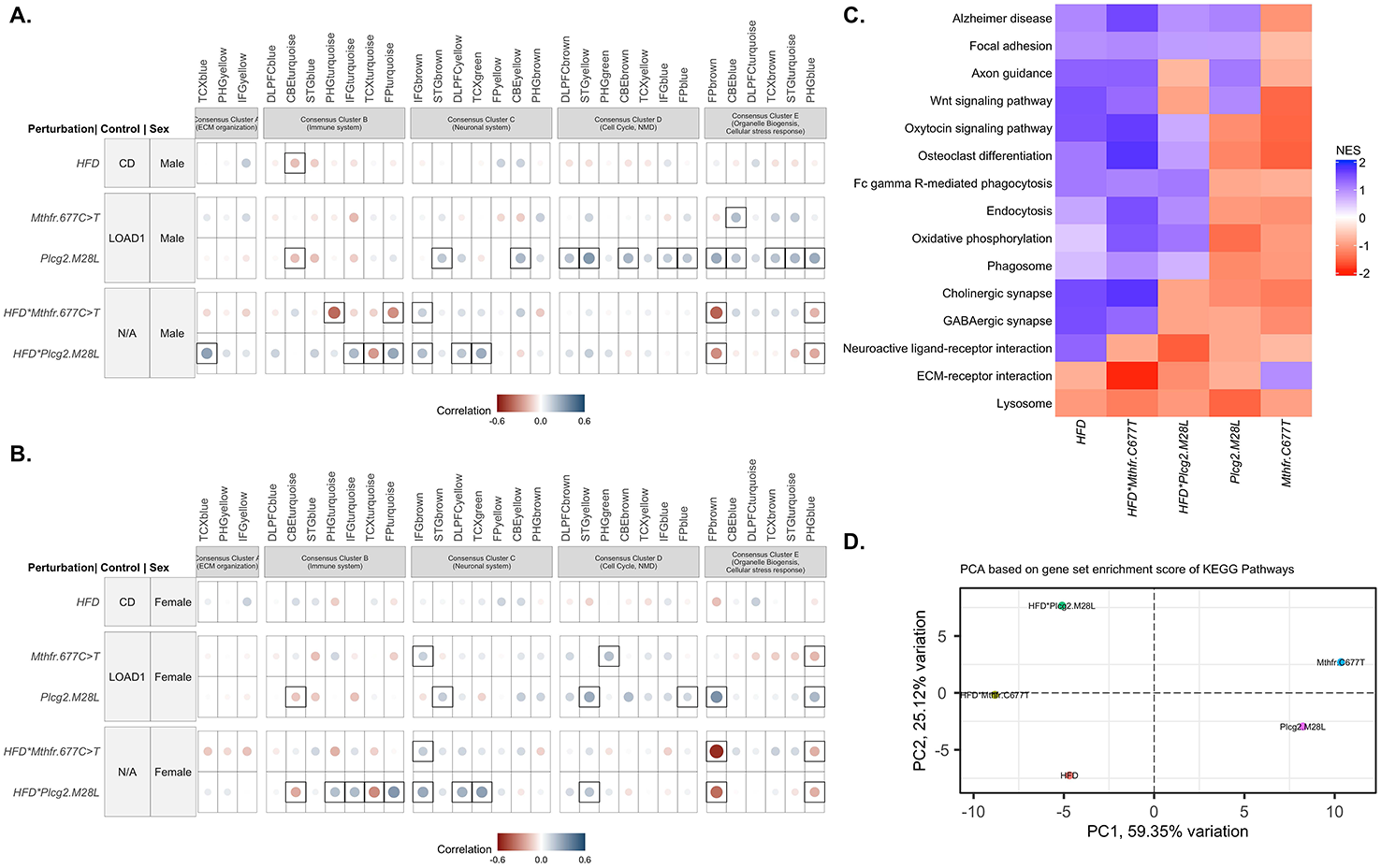

## Notes

### Competing Interest Statement

The authors have declared no competing interest.

